# AtRTD2: A Reference Transcript Dataset for accurate quantification of alternative splicing and expression changes in *Arabidopsis thaliana* RNA-seq data

**DOI:** 10.1101/051938

**Authors:** Runxuan Zhang, Cristiane P. G. Calixto, Yamile Marquez, Peter Venhuizen, Nikoleta A. Tzioutziou, Wenbin Guo, Mark Spensley, Nicolas Frei dit Frey, Heribert Hirt, Allan B. James, Hugh G. Nimmo, Andrea Barta, Maria Kalyna, John W. S. Brown

## Abstract

**Background:** Alternative splicing is the major post-transcriptional mechanism by which gene expression is regulated and affects a wide range of processes and responses in most eukaryotic organisms. RNA-sequencing (RNA-seq) can generate genome-wide quantification of individual transcript isoforms to identify changes in expression and alternative splicing. RNA-seq is an essential modern tool but its ability to accurately quantify transcript isoforms depends on the diversity, completeness and quality of the transcript information.

**Results:** We have developed a new Reference Transcript Dataset for Arabidopsis (AtRTD2) for RNA-seq analysis containing over 82k non-redundant transcripts, whereby 74,194 transcripts originate from 27,667 protein-coding genes. A total of 13,524 protein-coding genes have at least one alternatively spliced transcript in AtRTD2 such that about 60% of the 22,453 protein-coding, intron-containing genes in Arabidopsis undergo alternative splicing. More than 600 putative U12 introns were identified in more than 2,000 transcripts. AtRTD2 was generated from transcript assemblies of ca. 8.5 billion pairs of reads from 285 RNA-seq data sets obtained from 129 RNA-seq libraries and merged along with the previous version, AtRTD, and Araport11 transcript assemblies. AtRTD2 increases the diversity of transcripts and through application of stringent filters represents the most extensive and accurate transcript collection for Arabidopsis to date. We have demonstrated a generally good correlation of alternative splicing ratios from RNA-seq data analysed by Salmon and experimental data from high resolution RT-PCR. However, we have observed inaccurate quantification of transcript isoforms for genes with multiple transcripts which have variation in the lengths of their UTRs. This variation is not effectively corrected in RNA-seq analysis programmes and will therefore impact RNA-seq analyses generally. To address this, we have tested different genome-wide modifications of AtRTD2 to improve transcript quantification and alternative splicing analysis. As a result, we release AtRTD2-QUASI specifically for use in <u>Qu</u>antification of <u>A</u>lternatively <u>S</u>pliced <u>I</u>soforms and demonstrate that it out-performs other available transcriptomes for RNA-seq analysis.

**Conclusions:** We have generated a new transcriptome resource for RNA-seq analyses in Arabidopsis (AtRTD2) designed to address quantification of different isoforms and alternative splicing in gene expression studies. Experimental validation of alternative splicing changes identified inaccuracies in transcript quantification due to UTR length variation. To solve this problem, we also release a modified reference transcriptome, AtRTD2-QUASI for quantification of transcript isoforms, which shows high correlation with experimental data.

## Background

In plant and animal genomes, the majority of intron-containing genes undergo alternative splicing (AS). AS of precursor messenger RNAs (pre-mRNAs) can generate different transcript isoforms by selection of different splice sites [1-3]. A major consequence of AS is that mRNA AS variants are translated to produce different protein isoforms often with different or even antagonistic functions [1, 2]. In addition to increasing protein complexity, AS can regulate protein (and transcript) abundance by switching to production of transcript isoforms which are degraded by the nonsense-mediated decay (NMD) pathway [4-7].

In higher plants, AS is important in normal growth and development as well as in responses to biotic and abiotic stresses [8-13]. The significance of AS as a key regulator of gene expression is illustrated by 60-70% of intron-containing genes undergoing AS [14, 15]. Its functional importance is demonstrated in, for example, development, flowering time, the circadian clock, light signalling, seed dormancy, disease resistance and abiotic stress [16-28]. It is therefore essential that gene expression studies in plants take full account of the diversity of AS transcripts and assess the dynamic changes in expression at the individual transcript level to better understand how plant processes are controlled.

RNA-sequencing (RNA-seq) allows the assessment of differential expression of genes and transcripts through quantification of transcripts across a broad dynamic range. Nevertheless, accurate genome-wide identification and quantification of transcript isoforms from RNA-seq data remains a substantial challenge. With an available genome sequence the quantification of transcripts usually involves mapping of RNA-seq reads to the genomic reference and then construction of transcript isoforms as the first steps. Transcript expression levels are then inferred based on the number of aligned reads. Current analysis tools that identify and quantify transcripts from reads mapped to a genome include TopHat2/Cufflinks [29-32], RSEM [33, 34], eXpress [35], Bayesembler [36] and StringTie [37]. However, the determination of transcripts from short reads is inaccurate and generates incorrect, mis-assembled transcripts and misses *bona fide* transcripts that impacts the accuracy of transcript quantification [37-39]. For example, the assembly functions of two of the best performing programmes, Cufflinks and StringTie, generate 35-50% false positives, and quantification based on these transcript annotations often leads to inaccurate results [37]. In particular, for genes with multiple isoforms, the accuracy of transcript inference and quantification is poor [39]. Therefore, assembled transcripts and their quantification require rigorous experimental validation.

Rapid and accurate quantification of known transcripts can be achieved using programmes such as Sailfish and Salmon [40, 41] or Kallisto [42], which use lightweight algorithms to quantify the abundance of previously annotated RNA isoforms [40]. To analyse AS data in *Arabidopsis thaliana* we previously developed an Arabidopsis Reference Transcript Dataset (AtRTD) (now referred to as AtRTD1), and showed high correlation of AS using individual transcript abundances from Sailfish and Salmon and experimental data from high resolution (HR) RT-PCR [43]. AtRTD1 was made up of transcripts identified during RNA-seq analysis of a normalised library from 10-day old seedlings and flower tissue [14] and from transcripts in The Arabidopsis Information Resource version 10 (TAIR10) [44]. Here, we describe AtRTD2 where we have increased the number and diversity of Arabidopsis transcripts (over 82k unique transcripts from around 34k genes) by assembling ca. 8.5 billion 100 bp pairs of reads from RNA-seq libraries of a diverse range of plant material and incorporating transcripts from Araport11 Pre-release 3 [45, 46], hereafter Araport11. AtRTD2 contains higher confidence transcripts through the application of stringent filtering and quality control measures based on our knowledge of plant intron and splicing characteristics to reduce the number of probable false positive transcripts and other factors which could perturb quantification. Good correlation of quantification of AS using 127 AS events in 62 genes was observed between Salmon/AtRTD2 data and HR RT-PCR. However, for genes with transcripts which have variation in the lengths of their 5’ and 3’ untranslated regions (UTRs), quantification of transcript isoforms was often inaccurate. To reduce effects of this variation on the accuracy of quantification, we tested different modifications of AtRTD2 specifically for their ability to quantify AS accurately. As a result, we provide AtRTD2-QUASI (<u>Qu</u>antification of <u>A</u>lternatively <u>S</u>pliced <u>I</u>soforms) for use with programmes such as Sailfish, Salmon and Kallisto [40-42] and downstream AS analysis programmes such as SUPPA [47] to analyse or re-analyse RNA-seq data.

## Results

### Generation of AtRTD2

The main objective of generating AtRTD2 was to provide a new transcriptome for Arabidopsis for gene expression and, in particular, alternative splicing analysis. Well-annotated genomes and transcriptomes generally improve quantification of transcript isoforms, especially using RNA-seq analyses. We therefore wished to produce a reference transcript dataset that was as comprehensive and diverse as possible in terms of AS isoforms and contained the highest quality of supported transcripts. The quality of transcripts was achieved by applying stringent criteria to minimise the number of false, mis-assembled and poorly supported transcripts, and transcript fragments. To provide diversity of transcripts, two different extensive datasets of ca. 8.5 billion pairs of reads obtained from 285 RNA-seq runs of 129 libraries (Methods and Supplemental Methods: Table S1) were assembled into transcripts and merged with our previous AtRTD1 [43] and the recently released Araport11 transcript set [45, 46] (Figure 1). The two datasets were analysed separately in the different research groups and used different mapping programmes (STAR and TopHat2) but both used the transcriptome assembly functions in Cufflinks and StringTie and similar filtering approaches.

**Figure 1.**
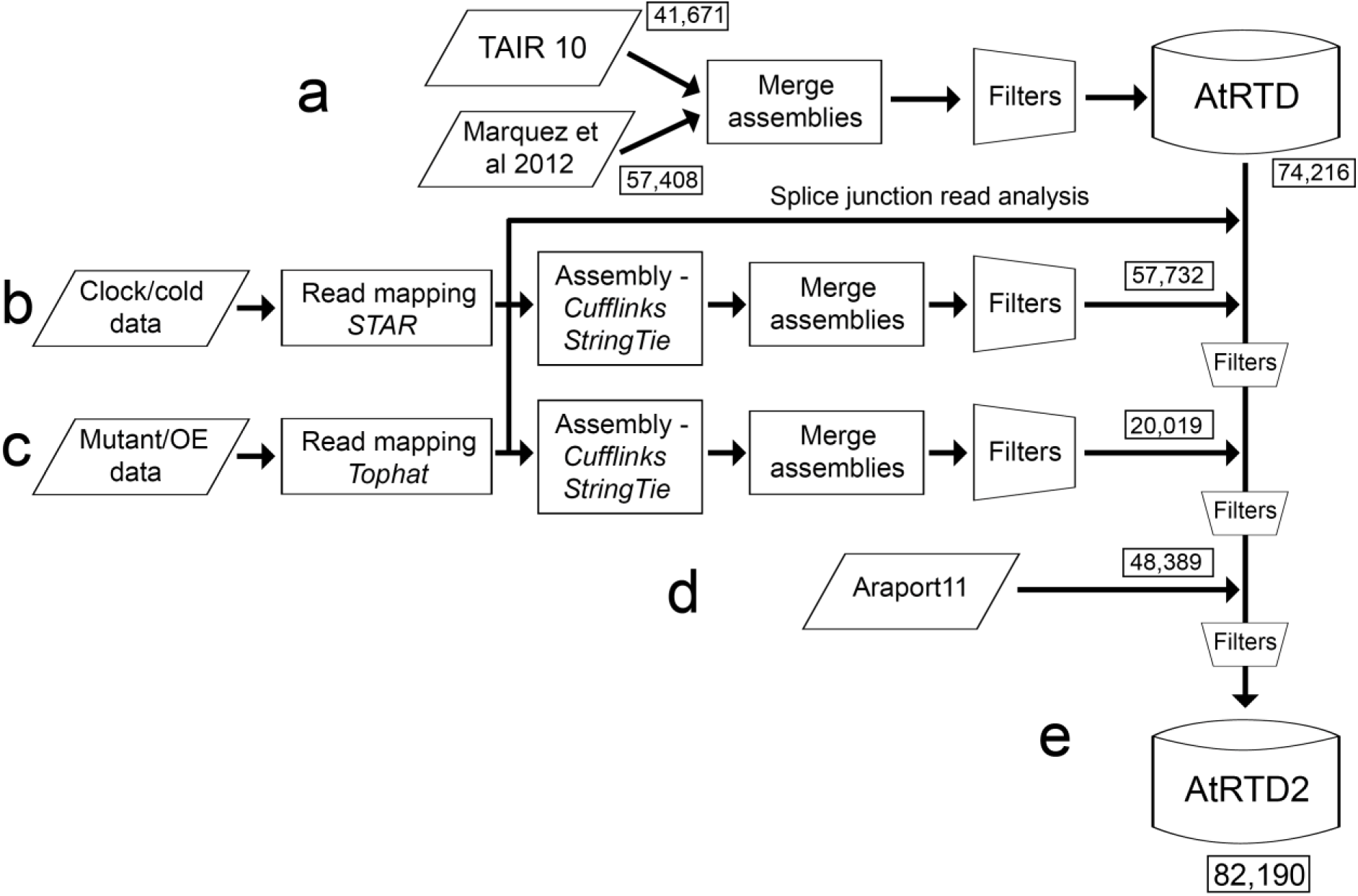
Construction of AtRTD2. **a**) The original AtRTD1 was generated from the merge of transcripts from TAIR 10 and from Marquez et al [14] and filtered to remove redundancy [43]. Using the splice junction data from the assemblies in **b**) and **c**), probable mis-assembled transcripts were removed; **b**) the dataset 1 RNA seq reads were mapped with STAR and then assembled using Cufflinks and StringTie. The assemblies were merged and filtered to give a transcriptome; **c**) the Dataset 2 RNA-seq reads were mapped with TopHat2, assembled, merged and filtered as above; **d**) Araport11 transcripts were then merged. Following each transcriptome merge, further filters (e.g. to remove redundancy) were applied (see Supplemental Figure 1); **e**) this generated the final AtRTD2.

To generate the AtRTD2 two different series of quality control filters were applied; the first at the transcript assembly stage and the second at the transcriptome merge stage (see Supplemental Figure S1). The assembly functions of Cufflinks and StringTie can generate large numbers of false positives [37, 38]. Following mapping of reads to the genome with STAR and TopHat2, we identified that both Cufflinks and StringTie generated novel splice junctions which were unsupported by reads. Therefore, the resulting transcriptome assemblies were initially filtered on the basis of splice junction quality. To this end, we combined the splice junction reads mapped by STAR and TopHat2 and kept only those with canonical splice sites (GT‥AG, GC‥AG and AT‥AC) which were present in at least ten reads in at least three sequencing samples/replicates (Dataset 1) or in three independent samples (Dataset 2), generating around 220k unique splice junctions. Applying these splice junctions to the transcript assemblies resulted in the removal of 49.0% and 28.3% of the transcripts from the initial Cufflinks and StringTie assemblies for Dataset 1 (see Supplemental Figure S1a). Similarly, using TopHat2/Cufflinks and TopHat2/StringTie on Dataset 2, 33.5% and 33.6% of assembled transcript models contained introns which were not supported by splice junction reads and were removed. Therefore, substantial numbers of transcripts assembled by both programmes here contained at least one intron with either non-canonical splice sites or with insufficient support by splice junction reads. The prediction of such “introns” by these programmes highlights the need to quality control assembled transcripts prior to quantification of transcript isoforms. Following the splice junction quality filter, antisense transcripts that were completely contained within an annotated gene in TAIR10 were removed along with transcripts from unknown genes and transcripts with no or very low expression (see, for example, Supplemental Figure S1a). The latter were removed on the basis that they likely represent mis-assembled transcripts. To this end, RNA-seq read data were analysed using Salmon, and only the transcripts which had a TPM >1 in at least three sequencing samples were kept. RNA-seq from Dataset 1 generated a transcriptome of 48,509 transcripts with Cufflinks and 50,942 with StringTie, of which 38,133 transcripts were identical (representing 78.6% and 74.9% of the Cufflinks and StringTie assemblies respectively) (see Supplemental Figure S2a). The remaining 10,376 and 12,809 transcripts were found in only the Cufflinks and StringTie transcriptomes, respectively (see Supplemental Figure S2a). For Dataset 2, the Cufflinks and StringTie assemblies contained 52,984 and 47,816 transcript models, respectively, of which 37,903 (71.5% for Cufflinks and 79.3% for StringTie) were identical between the two assemblies (see Supplemental Figure S2b).

To generate the final AtRTD2, the various total transcriptomes were merged together in a step-wise manner (Figure 1), and a second set of splicing and redundancy filters was applied after each merge (see Supplemental Methods and Supplemental Figure S1b). The application of the stringent filters will result in the loss of some transcripts with alternative transcription start sites or polyadenylation sites and of some *bona fide* transcripts which may be only expressed at very low levels or in a small number of specific cells. However, they have been removed currently for the purpose of generating a robust core dataset for transcript quantification and alternative splicing analyses in Arabidopsis. The AtRTD2 contains 82,190 transcripts where each transcript model represents a unique transcript isoform. Finally, to show that the introns in the AtRTD2 transcripts predicted by the splice junction reads were *bona fide* introns, we searched for sequence signatures of plant U2 and U12 introns using position weight matrices (PWMs) [14, 48]. The majority of introns (>99.8%) define typical plant introns with PWM values at both 5’ and 3’ splice sites of >60 (74.8% with PWM values >65 at each site) (data not shown). In addition, we identified 41 putative U12 AT‥AC and 589 putative U12 GU‥AG introns in 156 and 2,152 transcripts, respectively (PWM values of >75 and >65 at 5’ and 3’ splice sites, respectively).

### AtRTD2 has greater transcript diversity than AtRTD1

AtRTD2 was generated from assembly of two RNA-seq datasets (8.5 Bn paired end reads) and merger with transcript sets from Marquez et al [14], AtRTD1, TAIR10 and Araport11. Redundancy filters removed transcripts with the same intron co-ordinates which were shorter than other transcripts. The final AtRTD2 was made up of 33,674 unique transcripts from AtRTD1, of which 18,801 came from TAIR10 and 14,872 from Marquez et al. [14]; 7,518 and 10,460 new transcripts from Datasets 1 and 2, respectively, and 30,538 transcripts from Araport11 reflecting the extended 5’ and 3’ UTR sequences generated in Araport11 compared to TAIR10. The overall number of transcripts in AtRTD2 was increased by ∼18k transcripts in comparison to AtRTD1 (after removal of transcripts from AtRTD1 which were unsupported by splice junction reads – see Methods and Supplemental Methods). AtRTD2 has transcripts from over 34k genes which include non-coding (nc) RNA genes such as microRNA (miRNA), spliceosomal small nuclear RNA, small nucleolar RNA, transfer RNA genes etc. The majority of these genes do not undergo AS and will have a single transcript isoform. However, although some miRNA and long ncRNA precursors are alternatively spliced [49-51], our main focus is on alternatively spliced protein-coding genes. For the 27,667 protein-coding genes currently annotated in Araport11, AtRTD2 has a total of 74,197 transcript isoforms. Thus, there is an average of 2.68 transcripts per protein-coding gene in AtRTD2 reflecting higher transcript isoform complexity than found previously for all genes in TAIR10 (1.24), Marquez et al. [14] (2.40), and AtRTD1 (2.21) and for protein-coding genes in Araport 11 (1.75) (Table 1). The increased transcript complexity in AtRTD2 compared to TAIR10 and AtRTD1 is also shown by the increased number of genes with higher numbers of transcripts (see Supplemental Figure S3). A total of 13,524 protein-coding genes have at least one AS transcript in AtRTD2 (see Supplemental Figure S3) such that 60.23% of the 22,453 protein-coding, intron-containing genes in Arabidopsis undergo AS. Thus, AtRTD2 represents a non-redundant transcript dataset highly enriched in AS transcripts. Although different sources of transcripts have been used to generate the AtRTD2, all possible developmental stages and environmental conditions are not covered such that it is likely that some AS transcripts are missing. Therefore, new releases of AtRTD will be generated as other high quality RNA-seq data becomes available.

**Table 1.** Number of *Arabidopsis thaliana* genes and transcripts in different datasets. Number of transcripts and the average number of transcripts per gene are based on the total number of genes (TAIR10, AtRTD1 and AtRTD2) or the total number of genes detected [14] or on protein-coding genes (Araport11 and AtRTD2). ^a^ contains redundant transcripts which differ only by lengths of 5’ and 3’ UTRs ^b^ merged, non-redundant transcripts

|  | **Number of Genes** | **Number of Transcripts** | **Average Number of Transcripts per Gene** |
| --- | --- | --- | --- |
| **All genes** |  |  |  |
| TAIR10 | 33,602 | 41,671<sup>a</sup> | 1.24 |
| Marquez et al (2012) | 23,905 | 57,408 | 2.40 |
| AtRTD1 | 33,625 | 74,216<sup>b</sup> | 2.21 |
| AtRTD2 | 34,212 | 82,190 | 2.40 |
| **Protein-coding genes** |  |  |  |
| Araport11 | 27,667 | 48,389 | 1.75 |
| AtRTD2 | 27,667 | 74,194 | 2.68 |

<sup>b</sup> merged, non-redundant transcripts

### Quantification of transcripts with Salmon and AtRTD2

To demonstrate the utility of the AtRTD2, we have used it to quantify transcripts in RNA-seq data with the Salmon quantification tool [41]. To validate the quantification of the resulting changes in AS transcript ratios, we have compared the RNA-seq data to HR RT-PCR data generated using the same RNA samples. Previously, HR RT-PCR [52] proved to be instrumental for validating transcript assemblies from RNA-seq data: HR RT-PCR and Sanger sequencing of amplicons confirmed >90% of 586 assembled AS transcripts for 256 genes in the Arabidopsis normalized library RNA-seq [14]. Here, AtRTD2 transcript structures were compared to the amplicons in HR RT-PCR and the TPMs of individual transcripts used to calculate splicing ratios for each of the AS events or event combinations in that region (see Supplemental Figures S4, S7, S9-11). To directly compare the output from Salmon with HR RT-PCR, we analysed a total of 762 data points derived from 127 AS events from 62 genes and three biological replicates of the two time-points (T1 and T2, see Methods). In some cases, HR RT-PCR detected relatively low abundance AS transcripts which were not identified in RNA-seq and, therefore, to make direct comparisons, we calculated a splicing ratio by comparing the abundance of individual AS transcripts to that of the fully spliced (FS) transcript, which is usually the most abundant transcript and codes for the full-length protein. The Pearson’s correlation coefficient was 0.722 and the Spearman’s rank correlation coefficient was 0.804 for AtRTD2 (Figure 2a). These correlation coefficients were unexpectedly less than those observed with AtRTD1 – 0.905 and 0.907 respectively [43]. In general, the ratios using AtRTD2 tended to over-estimate the abundance of AS transcripts, and we observed that some genes with transcripts with variation in their 5’ and/or 3’ UTR lengths tended to show discrepancies between the splicing ratios obtained from HR RT-PCR and Salmon/AtRTD2.

**Figure 2.**
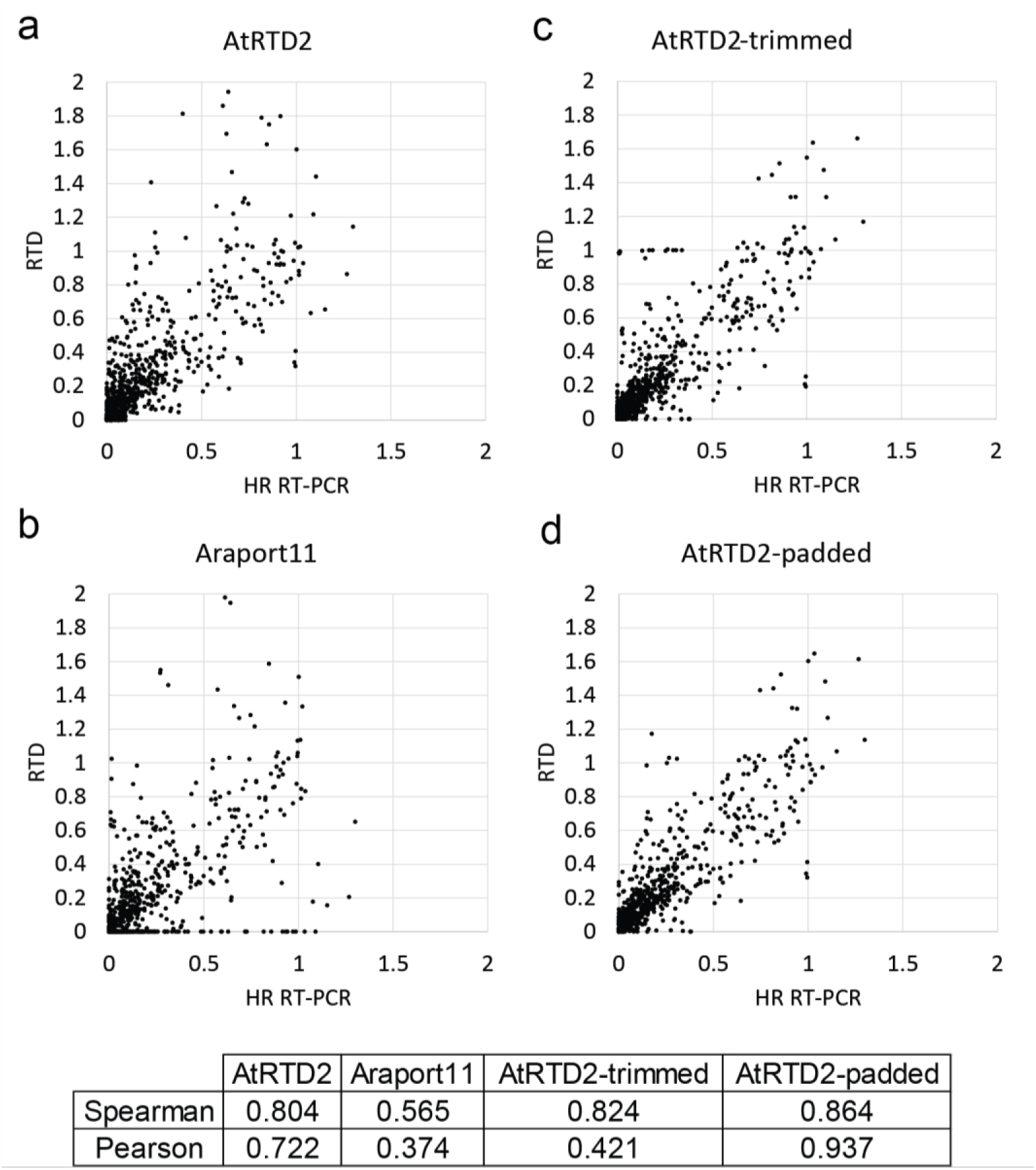
Correlation of splicing ratios calculated from the RNA-seq data and the High Resolution Reverse Transcription PCR (HR RT-PCR). Splicing ratios for 127 alternative splicing events from 62 *Arabidopsis thaliana* genes (three biological replicates of the time points T1 and T2) generated 762 data points in total. The splicing ratio of individual AS transcripts to the cognate fully spliced (FS) transcript was calculated from TPMs generated by Salmon and **a**) AtRTD2, **b**) Araport11, **c**) AtRTD2-trimmed, and **d**) AtRTD2-padded and compared to the ratio from HR RT-PCR. These AS/FS splicing ratios were calculated in this way to allow direct comparison with RNA-seq generated TPMs because for some genes, HR RT-PCR detected usually low level products representing AS events which were not present in the AtRTD2,. Correlation coefficients are given for each plot. Note that for clarity of the figures, 3, 7, 9 and 5 data-points with values that lie substantially outside the range of the graphs are not included in **a)-d)**, respectively, but are included in the correlations.

To show that the quality of AS change quantification depends on the completeness and diversity of transcript models in a reference transcriptome, we also compared the AS/FS splicing ratios obtained with HR RT-PCR to those derived from TPMs generated by analysing the same RNA-seq data using Araport11 as the reference transcriptome. We predicted that the smaller number of transcripts in Araport11 (ca. 48.4k transcripts) would impact transcript quantification and AS. We observed a Pearson’s correlation coefficient of 0.374 and a Spearman’s rank correlation coefficient of 0.565 (Figure 2b). The lower correlation with Araport11 reflects the fact that many AS events/transcripts are not present in the Araport11 transcriptome. At an individual gene level, the impact of missing transcripts is shown for *TRFL6* – AT1G72650 (Figure 3) and *FRS2* - AT2G32250 (see Supplemental Figure S4). In *TRFL6*, the transcript with retention of intron 4 makes up around 30% of expressed transcript isoforms but its absence in Araport11 affects the quantification of transcripts and thereby AS (Figure 3). In contrast, in *FRS2*, Araport11 uniquely provides a novel transcript (retention of intron 6) which represents around 25% of the transcript expression from this gene. Analysis of RNA-seq data using AtRTD2 minus Araport11 transcripts shows large changes in the relative abundances of the transcripts and in AS (see Supplemental Figure S4). Therefore, it is clear that the more comprehensive a reference transcriptome is, the better will be the accuracy of measuring AS isoforms and their contribution to gene expression.

**Figure 3.**
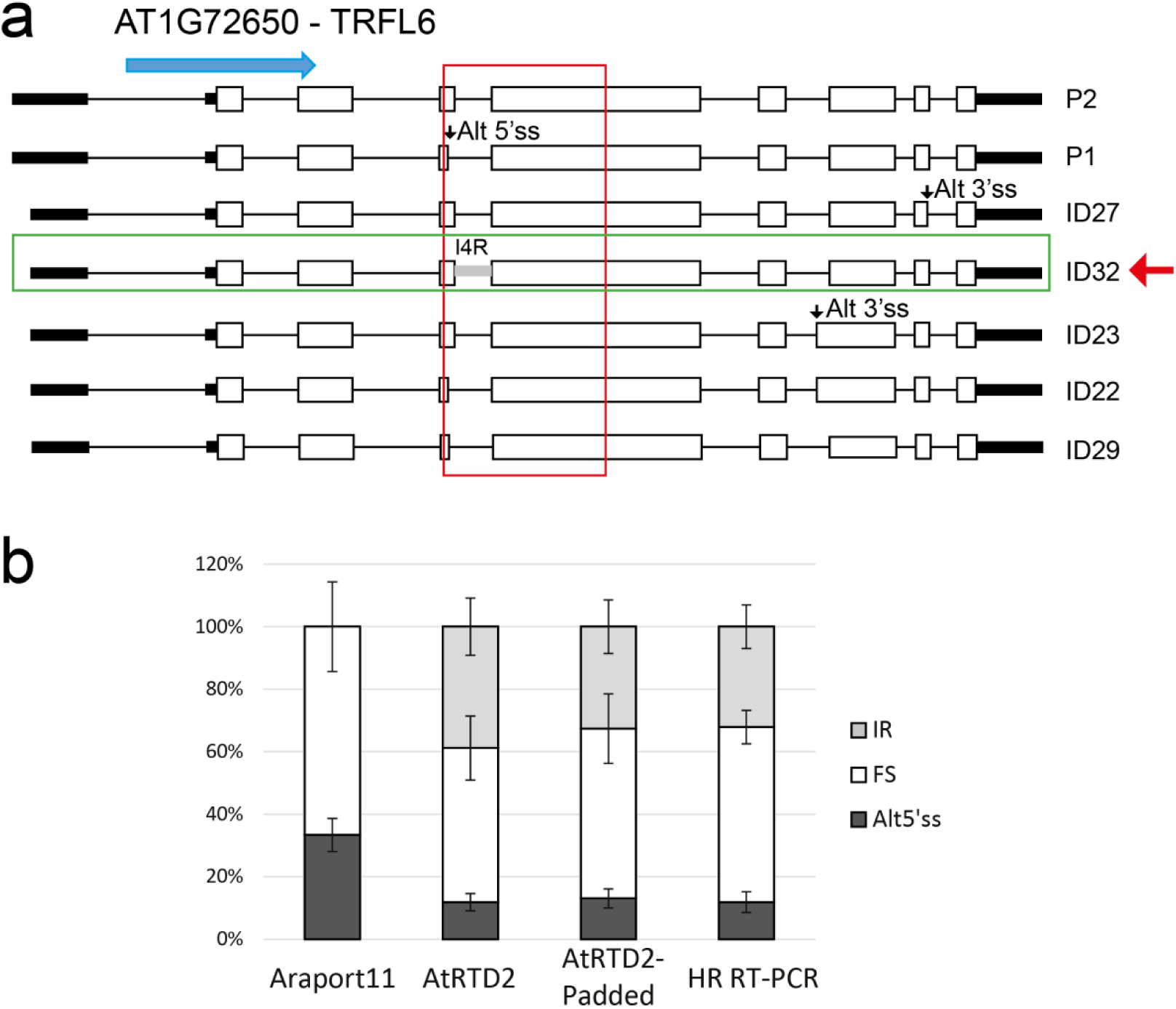
Missing reference transcripts affect transcript and AS quantification. **a**) AT1G72650 (*TRFL6*) has four different alternative splicing events (an Alt5’ss in exon 4; two Alt3’ss in exons 7 and 8, and intron retention of intron 4 (I4R) contained in seven transcripts. HR RT-PCR across exons 4 and 5 detects the fully spliced product, Alt5’ss and I4R (red box). Blue arrow – direction of transcription. **b**) AS splicing ratios calculated from TPMs of all seven transcripts from analysis of RNA-seq data (T1) using Araport11, AtRTD2 and AtRTD2-padded (see main text) as reference transcriptomes and compared to HR RT-PCR (T1 of dataset 1). The absence of a transcript with I4R in Araport11 (ID32 in AtRTD2 – green box) affects the quantification of transcripts and AS.

### Variation in UTR length among transcripts affects accuracy of quantification of isoforms and AS events

To address whether variation in UTRs affected quantification of transcript isoforms, we investigated the genes/AS events used in the HR RT-PCR analysis. Such UTR variation may be due to genes which have *bona fide* alternative transcription start sites or polyadenylation sites. Alternatively, they may reflect stochastic transcription start variation [53], artefacts of reverse transcription or internal priming in cDNA/EST cloning and sequencing or in library preparation for RNA-seq (e.g. in TAIR10; [54]), variation due to *in vivo* or *in vitro* RNA degradation or differences in transcript assembly. We made the assumption that much of the variation in 5’ and 3’ UTR length of transcripts is likely to be due to transcripts being incomplete and missing terminal regions (i.e. not full-length) or to local variation or heterogeneity in transcription starts [53] or cleavage/polyadenylation sites [54].

We reasoned that modifying transcripts so they had the same start and end co-ordinates should improve quantification of isoforms. Therefore, to demonstrate that UTR length variation impacted quantification of such genes, we took two approaches: 1) we trimme d transcripts from the 5’ and 3’ ends to the co-ordinates of the transcript(s) that covered the smallest region of the gene (see Supplemental Figure S5a and S5b) or 2) we padded genomic sequence from the ends of the shorter transcripts up to the co-ordinates of the end of the transcript(s) that covered the biggest region of the gene (see Supplemental Figure S6a and S6b). These modifications were performed genome-wide on AtRTD2 to generate AtRTD2-trimmed and AtRTD2-padded, and the RNA-seq data were analysed with Salmon and the modified AtRTD2 datasets. Splicing ratios were again calculated and compared to HR RT-PCR for the 127 AS transcripts. Both the trimmed and padded versions of AtRTD2 improved the Spearman’s rank correlation (0.824 and 0.864 respectively) compared to AtRTD2 (Figure 2). The Pearson’s correlation also improved with the AtRTD2-padded version (0.937) but was greatly reduced for the trimmed version (0.421). Although trimming of 5’ and 3’ ends for many genes resulted in a higher correlation with HR RT-PCR, correlations for some genes were markedly different. By examining the effects of trimming in detail for these genes, we identified cases where trimming gave rise to new UTR sequence length variation affecting quantification (see Supplemental Figure S5c and S5d). For example, when the 5’ end of a shorter transcript corresponded to a position in a 5’ UTR intron (see Supplemental Figure S5c), trimming to this position often resulted in the longer transcripts losing exon sequences and becoming shorter, thereby introducing new UTR variation (see Supplemental Figure S5d). The impact on isoform quantification is illustrated for AT3G23280 (*XBAT35*) where trimming to the length of the shortest transcript (AT3G23280.s1) removes exons 1 and 2 of the other transcripts, generating new UTR variation and causing large changes in the TPM values of the transcripts (see Supplemental Figure S7).

We found an overall increased correlation of AS/FS ratios from HR RT-PCR with the AtRTD2-padded version (Figure 2). To examine whether the effect was due to quantification using Salmon, we also analysed our RNA-seq data with Kallisto using both AtRTD2 and AtRTD2-padded. We found that Kallisto generated very similar results and also showed an improved correlation of HR RT-PCR data with the AtRTD2-padded version (see Supplemental Figure S8). We examined the effects of padding of AtRTD2 in detail for some genes/transcripts. TPM values were similar using AtRTD2 and AtRTD2-padded for genes where transcripts had little or no differences in their 5’ or 3’ ends (for example see AT5G05550 (*VFP5*) - Supplemental Figure S9). For genes with 5’ and 3’ end variation in the first and/or last exon, AtRTD2-padded gave more accurate TPM values. This was clearly shown by the three-way corroboration of AS/FS values from HR RT-PCR, from TPMs from analysing the RNA-seq data with Salmon and Kallisto using AtRTD2-padded, and from manual counting of splice junction reads in a read alignment viewer. For example, there is a 10-fold difference in the AS/FS splicing ratios of the Alt3’ss event in *CRY2* between the HR RT-PCR data, Salmon/AtRTD2-padded and read counts when compared to analysis with AtRTD2 or Araport11 (Figure 4). Further examples are shown for genes encoding a RING/U-box superfamily protein and *HSF3* (see Supplemental Figures S10 and S11). While trimming involving an intron (e.g. in the 5’ UTR) had negative effects on accuracy of transcript quantification (Figures S5, S7a and S7b), padding of a shorter transcript which ended within an intron did not appear to affect quantification greatly. For example, a transcript ending within a 5’ UTR intron is indicative of an intron retention event and padding effectively generates the full intron retention (Figures S6c and S6d). This is illustrated by AT4G35800 which has an intron in the 3’ UTR (intron 13) and a transcript which terminates in the intron. Padding generates a transcript with retention of intron 13 which has been shown to occur and to be up-regulated in the cold [55] (data not shown).

**Figure 4.**
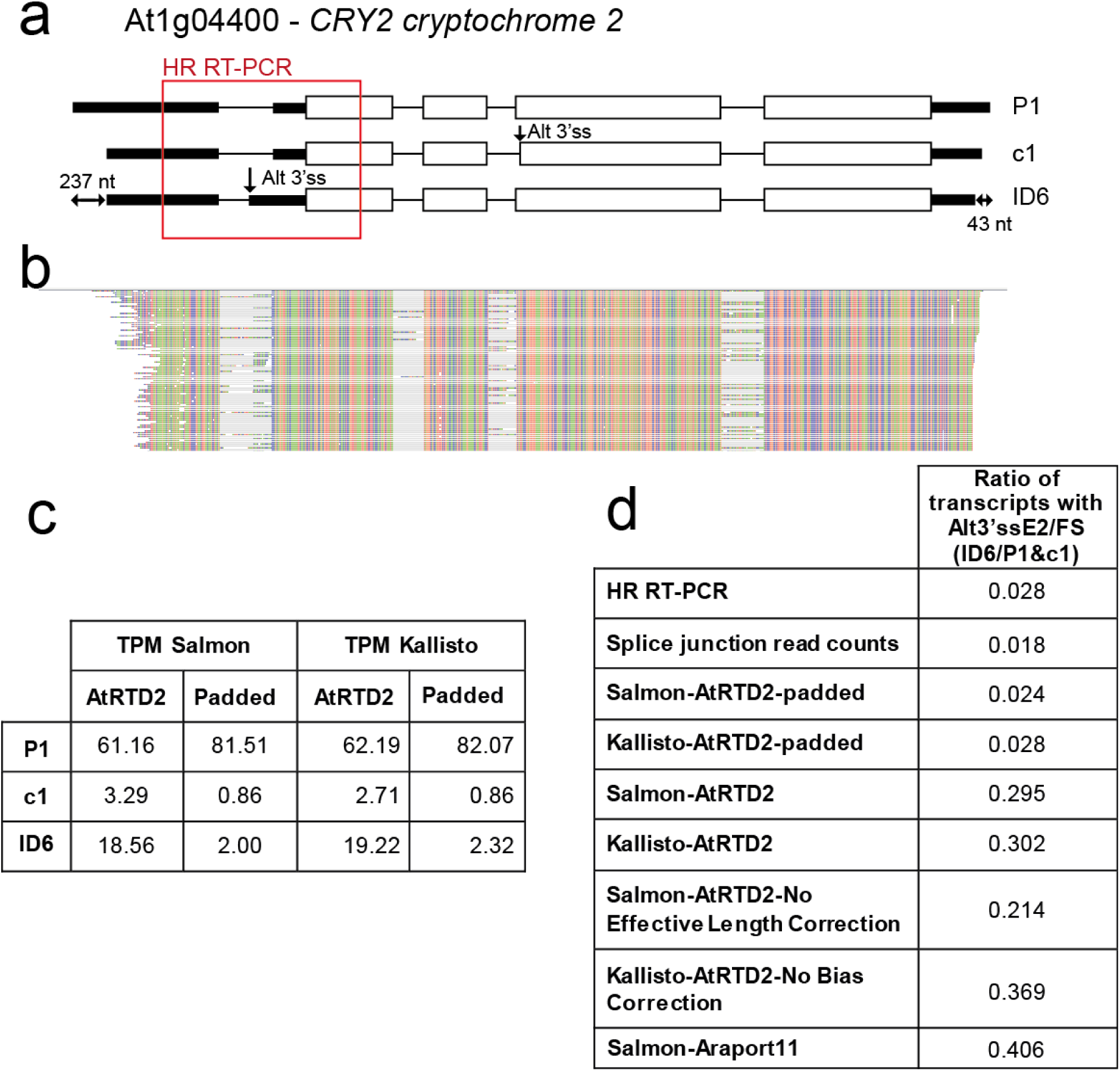
Variation in UTR lengths affects transcript and AS quantification. **a**) AT1G04400 (*CRY2; CRYPTOCHROME 2*) has two different AS events (an Alt3’ss exon 2 and Alt3’ss exon 4). The difference between shortest (ID6) and longest (P1) transcripts is 237 and 43 nt at the 5’ UTR and 3’ UTR, respectively. **b**) Read alignment showing the relatively low level of Alt3’ss splice junction reads. **c**) Average TPMs of the 3 transcripts using Salmon and Kallisto and the AtRTD2 and AtRTD2-padded; **d**) Splicing ratios of the Alt3’ss in exon 2 to FS (fully spliced) transcripts with HR RT-PCR, manual counting of splice junction reads in this region using a read alignment viewer [61], Salmon and Kallisto with AtRTD2-padded and AtRTD2, Salmon and Kallisto with uncorrected functions with AtRTD2, and Salmon with Araport11.

To elucidate the effect of 5’ and 3’ UTR variation on accuracy of transcript quantification among all of the genes/transcripts analysed by HR RT-PCR, we measured the length of padding for each gene by calculating the difference between the 5’ and 3’ ends of each transcript and shortest transcript of the gene and then adding the 5’ UTR and 3’ UTR variation together (see Supplemental Figure S12a and S12b). We divided these into three size classes: 0-100 nt, 101-600 nt and >600 nt and compared the correlation between HR RT-PCR and AtRTD2 or AtRTD2-padded (see Supplemental Figure S12c). In particular, we observe greatly improved correlation for transcripts where padding has added between 100 and 600 nt at the 5’ and 3’ ends. When the UTR variation is >600 nt, there is still an improvement in the correlation with padding but the underlying correlation with AtRTD2 is poor. Thus, variation at the 5’ and 3’ UTRs among transcripts from the same gene can greatly affect accuracy of transcript quantification. We therefore suggest that AtRTD2-padded is a useful tool in quantification of AS transcript isoforms for many genes and release it here as AtRTD2-QUASI specifically for use in <u>Qu</u>antification of <u>A</u>lternatively <u>S</u>pliced <u>I</u>soforms. AtRTD2-QUASI overcomes problems of local variation and heterogeneity in the 5’ and 3’ ends of transcripts. The caveat to its utility is that, because of the assumption on which it is based, it may not be appropriate for quantification of transcripts with *bona fide* alternative transcription start sites or functionally different polyadenylation sites.

Finally, the AtRTD2-QUASI modified reference set improves the accuracy of isoform quantification and thereby differential expression analyses. Why artificially extended shorter transcripts delivers this increased accuracy is unclear. Most transcript quantification and differential expression programmes have algorithms to correct for read distribution variation based on the effective length of transcripts and for bias of read distribution towards the ends of transcripts. The effective length of a transcript is defined as the convolution of fragment length distribution and transcript length+1 [29, 30], which essentially represents the positions in the transcript that can generate a valid fragment. Its value is equal to the transcript length+1 minus the mean fragment length. Given that the mean fragment length is the same for an experiment, the effective length correction is a universal shift of length for all transcripts. The differences of start and end site of the transcripts, due to incompleteness or false extension of the transcripts of a gene, are not accounted for by effective length correction. Fragment bias is characterized in two ways [56]: 1) positional bias, which describes a local effect that fragments are preferentially located towards either the beginning or end of the transcripts, and 2) sequence-specific bias, which is a global effect where some nucleotides at the beginning or end of transcripts affect their chances of being selected for sequencing. The second aspect is unlikely to be affected by the different length of 5’ and 3’ UTRs in the transcript annotations. However the positional bias is a statistical measure that is estimated based on all transcripts, or a few transcript bins segregated by transcript length [56]. For isoforms with different lengths of 3’ or 5’ UTRs, the positional bias at the same corresponding genomic location within the isoforms, which should be the same, could be characterised and quantified very differently such that, again, differences in 3’ and 5’ UTRs due to incompleteness or false extension are not well accounted for by the positional bias corrections. We therefore tested these functions to see their effect on the improved accuracy obtained with padding by analysing the RNA-seq data using Salmon with the EffectiveLengthCorrection function turned off and using Kallisto with the read bias correction turned off using both AtRTD2 and AtRTD2-QUASI as references. The values of TPMs and of AS/FS splicing ratios were similar to the inaccurate results obtained with AtRTD2 when these corrections in operation (see Figure 4 and Supplemental Figures S10 and S11). This suggests that the phenomenon that we have observed has a different basis. Examination of the transcript structures in the examples used here identifies three features which may be responsible for the less accurate quantification of transcripts when transcripts show 5’ and/or 3’ UTR variation: longer transcripts containing unique sequences, the different degrees of overlap among multiple transcripts or the inconsistency between read coverage from the experimental data and the longer transcripts from the AtRTD (see Supplemental Figure S13).

### Translation of AtRTD2 for proteomic analyses

An important question in alternative splicing is to what extent it generates proteome complexity. To date, this has been difficult to assess because of our relatively limited knowledge of AS isoforms. AtRTD2 contains a diverse set of supported transcripts. We therefore characterised the coding capacity of the AtRTD2 transcripts. Instead of translating the transcripts in all six reading frames to derive peptide sequences, we used an algorithm which fixed the translation start site AUG for each gene on the basis of the annotated reference gene model in TAIR10 such that translations terminated at the first stop codon from the AUG. This process thereby takes into account the real coding capacity of different transcript isoforms such as transcripts containing premature termination codons (PTCs) or changes of open reading frame (ORF) caused by AS [57]. *In silico* trypsin digestions generated peptides from each transcript and identified 399,701 peptides of which 200,867 are unique to specific AS isoforms.

## Discussion

In this paper we report a new *Arabidopsis thaliana* Reference Transcript Dataset (AtRTD2) for use in quantification of individual transcript abundances using the lightweight alignment programmes such as Salmon and Kallisto. The new AtRTD2 release contains around 82k non-redundant transcript isoforms where each transcript from a gene represents a different alternatively spliced transcript with at least one different splicing event. In addition to increased diversity of isoforms, stringent criteria have been applied to support the AtRTD2 transcripts. The AtRTD2 resource contains a significantly higher number of transcript isoforms than other current collections such as TAIR10 and Araport11. Accurate quantification of transcripts is essential for downstream differential expression analyses and depends on the diversity, quality and completeness of the reference transcripts (missing and mis-assembled can severely affect quantification) and thereby the transcriptome. Here, we have shown the drastic effects on quantification of missing transcripts both at the individual gene and transcriptome reference levels. Although AtRTD2 is likely to still be incomplete, further short and long read data can be continually incorporated to eventually generate a full *A. thaliana* transcriptome. Finally, extensive validation of transcripts and their abundance is important in assessing the quality of RNA-seq data and its analysis. Previously, we developed HR RT-PCR for analysis of changes in alternative splicing [5, 19, 24, 52, 58, 59] and have used it to validate transcript assemblies from RNA-seq data [14]. Here, we have performed extensive validation with HR RT-PCR and showed good correlation between splicing ratios calculated from TPMs from RNA-seq data analysed with both Salmon and Kallisto and the HR RT-PCR. However, this detailed analysis also identified inaccurate quantification for genes with variation in the lengths of their 5’ and 3’ UTRs and we have, therefore, generated another resource, AtRTD2-QUASI, which greatly improves the accuracy of quantification for many genes (see below).

RNA-seq is now widely used in transcriptome/expression analyses in plants to examine a wide range of developmental and environmental questions. In Arabidopsis, most RNA-seq data has been analysed with the TopHat/Cufflinks/Cuffdiff pipeline using TAIR9 or TAIR10 as references, despite the knowledge that these datasets are relatively poor in their coverage of AS transcripts. In addition, recent comparisons of different assembly and quantification programmes have pointed to the high degree of inaccuracy in transcript assembly of the most widely used programmes [37, 38]. There is wide variation in the performance of different computational methods for detection of plant AS in RNA-seq data, and the quality of annotation can have a major impact [60]. RNA-seq analysis using the combination of the AtRTD2 transcriptome with Salmon and extensive validation has allowed us to identify a major issue in the accuracy of quantification due to transcripts from the same gene having different lengths of 5’ and/or 3’ UTRs and that correction functions within quantification programmes do not appear to deal effectively with this problem. Thus, current methods of RNA-seq analysis in Arabidopsis are likely to be inaccurate at a number of different levels which represents a major problem for plant RNA-seq given that Arabidopsis has the most advanced genome and transcriptome annotation.

The effects of transcript length variation (edge bias) have been observed for some human transcripts in quantification of AS [47]. For example, when different transcripts of the same gene contain or do not contain UTR sequences, reads from the UTRs are disproportionately assigned to the transcript containing the UTR sequence and therefore can greatly affect the accuracy of quantification [47]. Here, we observe an opposite effect where variation in 5’ or 3’ UTRs is often associated with few or no reads and accuracy of quantification of isoforms is affected. There are a number of possible sources of such UTR variation. Firstly, some will reflect the use of *bona fide* alternative transcription start sites (TSS). Methods such as 5’ RACE have demonstrated alternative TSS for a small number of genes but genome-wide information is not extensive. Transcription can be stochastic, often starting in a region of the promoter with no one single nucleotide being the transcription start site such that transcripts may already have variation in the 5’ UTR [53]. Variation in the 3’ UTR can be generated by alternative polyadenylation. In Arabidopsis, genome-wide information is available from direct RNA sequencing and ca. 75% of genes have more than one polyadenylation site but most reads were associated with a preferred site [54]. Indeed, variation in polyA sites is often associated with multiple overlapping polyA signals perhaps ensuring termination in the compact genome [54]. Legacy transcripts, such as those in TAIR10, are derived from cDNA/EST cloning and sequencing and some contain no annotated UTRs, beginning with the translation start site and ending with the stop codon, although much of this latter variation has been re-annotated [54] and incorporated in the Araport11 transcripts. Variation of a few to tens of nucleotides may arise from artefacts of reverse transcription, priming of oligo-dT to internal sequences such that full-length cDNAs are not produced or different protocols of cDNA library preparation prior to RNA-seq. Finally, variation may be due to *in vivo* or *in vitro* RNA degradation.

As much of the variation in UTRs is likely to be due to uncertainty of the 5’ and 3’ ends of a transcript and as the discrepancy in quantification using AtRTD2 suggests that many of the most widely used RNA-seq analysis programmes are unable to take UTR length variation into consideration effectively, we have also developed AtRTD2-QUASI for accurate quantification of AS isoforms. We propose that RNA-seq data is analysed with Salmon or Kallisto using AtRTD2-QUASI because quantification data agrees well with data obtained experimentally (HR RT-PCR) and data derived from manually counting splice junction reads. The three-wa y corroboration of results suggests that AtRTD2-QUASI is a practical solution to obtaining good quantitative data on transcript isoforms. Although AtRTD2-QUASI improves the quantification markedly for the majority of genes, particular gene arrangements may not be reported on accurately and data should be treated with caution. For example, if quantification of genes with different *bona fide* transcription start sites and with different polyadenylation sites is performed by Salmon/Kallisto with AtRTD2-QUASI, experimental data will be required to decide on which output to use or whether either is appropriate.

Finally, we have translated the >82k unique transcripts from the authentic translational start site to reflect the most realistic open reading frames in different AS isoforms (e.g. AS events that are in frame, change frame or generate PTCs). This approach differs from the method of translating transcript isoforms in TAIR, based on the longest ORF which often results in the depiction of aberrant ORFs beginning at an AUG downstream of the authentic translation start site, instead of depicting the presence of PTC(s) [57]. Fixing the translational start site provides the means to identify peptides which are unique to specific AS isoforms which will benefit proteomic analyses.

## Conclusions

In this study we have developed a new Reference Transcript Dataset for *A. thaliana* (AtRTD2) for quantification of transcripts and changes in alternative splicing. We have demonstrated the impact that incomplete transcriptomes can have on quantification and that, therefore, transcriptomes need to be constantly refined with new data. Our analysis has also demonstrated the value of extensive experimental validation of RNA-seq data. In addition, we found that variation in the lengths of 5’ and 3’ UTRs of transcripts from the same genes perturbed accurate transcript quantification, and that current programmes appear unable to correct for such variation. This is likely to impact RNA-seq analyses of all organisms. We have generated AtRTD2-QUASI as a practical solution for quantifying transcript splice isoforms in Arabidopsis.

## Methods

### Plant material for RNA-seq and datasets

Two different extensive RNA-seq datasets were generated for a range of diverse genetic lines and treatments in the *Arabidopsis thaliana* Col-0 background (see Supplemental Methods: Table S1). These two combined datasets included 285 RNA-seq runs obtained from 129 libraries. Dataset 1 was from samples of a time-course of adult Arabidopsis plants (5 weeks old) transferred from 20°C to 4°C (unpublished data). Plants were sampled every three hours for the 24 h period at 20°C directly before transfer to 4°C, and for the first and fourth days following transfer (26 time-points in 78 libraries - a total of 234 total biological/sequencing repeats) (see Supplemental Methods: Table S1). Dataset 2 consisted of RNA-seq data from 51 libraries generated from samples of over-expression and mutant lines, some of which were treated with flagellin 22 (flg22) or mock-treated with water (unpublished data) (see Supplemental Methods: Table S1). The genetic lines were over-expression lines and knockout mutants of the serine-arginine-rich (SR) splicing factor genes, At-*RS31* (AT3G61860) and At-*RS2Z33* (AT2G37340); the mutant of the DNA cytosine methyltransferase MET-1 (AT5G49160), *met1-3*; and mutants of three MAP kinase genes (AT3G45640, AT4G01370 and AT2G43790), *mpk3*, *mpk4* and *mpk6*. The latter were treated with flg22 or mock-treated, and wild type Col-0 controls were included for all of the above. Dataset 2 also included transcripts from TAIR10 and from RNA-seq from a normalised library of wild-type flowers and 10 day old seedlings [14] used in the construction of AtRTD1 [43]. The total number of 100 bp paired-end reads generated in the two datasets of RNA-seq was 4.76 and 3.73 Bn pairs of reads, respectively, such that a total of ca. 8.5 Bn pairs of reads (17 Bn paired-end reads) entered the assembly pipeline.

### Generation of AtRTD2

A detailed description of the transcript assembly and parameters used, merging with the original AtRTD1 [43] and Araport11 is given in Supplemental Methods and shown schematically in Figure 1. Briefly, RNA-seq reads from Datasets 1 and 2 were mapped to the genome using STAR and TopHat2 respectively, and transcripts for both datasets were assembled with both Cufflinks and StringTie. Transcripts supported by non-canonical junctions or low abundance splice junction reads were removed. Transcripts from unknown genes, antisense transcripts and low abundance transcripts were also filtered out. The resulting Cufflinks and StringTie transcriptome assemblies were merged and redundant transcripts were removed. AtRTD1 was re-assessed using the splice junction sequence set generated here and 10,397 transcripts deriving from Marquez et al. [14] were removed. The modified AtRTD1 was then merged with the Dataset 1 transcriptome and then with that of Dataset 2, with a series of quality filters being applied at each step (see Supplemental Methods). Finally, the resulting transcriptome was merged with Araport11 transcript assemblies [45, 46] and filtered again to give the new Arabidopsis transcriptome, AtRTD2.

### Experimental validation of the quantification of splicing ratios by High Resolution RT-PCR

To validate the quantification of splicing ratios from transcript isoforms, HR RT-PCR [52] was performed on RNA from two different time-points (each with three biological repeats): dawn at 20°C (T1) and in the middle of the dark period four days after transfer to the low temperature (T2). T1 and T2 are from the same plant material as used for RNA-seq of Dataset 1. A total of 762 data points (127 AS events from 62 genes and three biological replicates of the two time-points) were analysed by HR RT-PCR using gene-specific primers (Table S2). Primer pairs covering the AS events in these genes, where the upstream primer was end-labelled with a fluorescent tag, were used in RT-PCR reactions with 24 cycles of PCR as described previously [5, 14, 24, 52] and separated on an ABI 3730 automatic DNA sequencing machine. The abundance of RT-PCR products was analysed with GeneMapper software and splicing ratios were calculated from peak areas of each product. We analysed the RNA-seq data from the same regions of these 62 genes and used the transcripts per million (TPM) values of the transcripts generated by Salmon on the RNA-seq data to calculate splicing ratios. For comparison with HR RT-PCR, Spearman and Pearson correlations were computed on splicing ratios.

### Modification of AtRTD2 for quantification of transcript abundances and generation of AtRTD2-QUASI

The AtRTD2 was modified to examine the effects of transcript 5’ and/or 3’ end length variation in genes on isoform quantification. Transcripts were trimmed at the 5’ end and 3’ ends. Trimming was to the end co-ordinates of the transcript that covered the shortest region of the gene and was achieved using in-house scripts (see Supplemental Figure S5). Alternatively, transcripts in AtRTD2 were padded to give the transcripts of each gene the same 5’ and 3’ ends. Shorter transcripts were extended to the co-ordinates of the transcript that covered the longest region on the gene by adding the cognate genomic sequence by in-house scripts (see Supplemental Figure S6). The AtRTD2-padded version was called AtRTD2-QUASI.

### Translation of AtRTD2 and new peptide database

The main consequences of AS are either to alter the protein-coding sequence to generate protein variants or introduce premature stop codons/long faux 3’UTRs which can target transcripts to the NMD pathway. In order to assess the consequences of an AS event(s), translation of the transcript from the authentic translation start site is required [57]. We therefore developed an algorithm which defined the position of the translation start AUG in the transcripts of a gene (usually that of the TAIR reference gene model) and used this translation start site as the reference point for translation of all of the transcripts. This generated realistic translations which were used in *in silico* trypsin digestions to generate a peptide database consisting of nearly 400k peptides of which half could be assigned to specific AS isoforms.

## Availability of data and materials

The datasets supporting the conclusions of this article are available in the James Hutton Institute repository, [ics.hutton.ac.uk/atRTD/]. Datasets are: AtRTD2, AtRTD2-QUASI and can now be used to analyse or re-analyse Arabidopsis RNA-seq data.

## Supplemental Files

**Supplemental Methods:** File contains supplemental methods.

**Supplemental Figures:** File contains supplemental figures.

## Authors contributions

AJ, CC, NF, NT, YM harvested material and extracted RNA; RZ, YM, PV carried out RNA-seq transcript assemblies and data analyses; RZ, CC, JB, NT, WG, MK participated in RNA-seq data analyses, interpretation and generation and analysis of modified AtRTD2; CC, NT performed the HR RT-PCR and analyses; MS developed the translation tool and performed the translation analyses; YM carried out the PWM analyses and generated the peptide database; JB, CC, RZ drafted the manuscript and all authors participated in its correction and have read and approved the final manuscript; AB, JB, HH, HN, MK acquired the funding.

## Acknowledgements

We thank Janet Laird (University of Glasgow) and Linda Milne, Micha Bayer and Iain Milne (James Hutton Institute) for technical assistance.

